# An individual differences approach to semantic cognition: Divergent effects of age on representation, retrieval and selection

**DOI:** 10.1101/182170

**Authors:** Paul Hoffman

## Abstract

Semantic cognition refers to the appropriate use of acquired knowledge about the world. This requires representation of knowledge as well as control processes which ensure that currently-relevant aspects of knowledge are retrieved and selected. Although these abilities can be impaired selectively following brain damage, the relationship between them in healthy individuals is unclear. It is also commonly assumed that semantic cognition is preserved in later life, because older people have greater reserves of knowledge. However, this claim overlooks the possibility of decline in semantic control processes. Here, semantic cognition was assessed in 100 young and older adults. Despite having a broader knowledge base, older people showed specific impairments in semantic control, performing more poorly than young people when selecting among competing semantic representations. Conversely, they showed preserved controlled retrieval of less salient information from the semantic store. Breadth of semantic knowledge was positively correlated with controlled retrieval but was unrelated to semantic selection ability, which was instead correlated with non-semantic executive function. These findings indicate that three distinct elements contribute to semantic cognition: semantic representations that accumulate throughout the lifespan, processes for controlled retrieval of less salient semantic information, which appear age-invariant, and mechanisms for selecting task-relevant aspects of semantic knowledge, which decline with age and may relate more closely to domain-general executive control.

## Introduction

Semantic knowledge, of the meanings of words and properties of objects, shapes our understanding of the world and guides our behaviour. Each of us has a wide range of conceptual knowledge distilled from a lifetime of experiences. Effective semantic cognition – that is, the ability to use semantic knowledge to complete cognitive tasks – requires us to both *represent* this information in an accessible form and to *control* how we access and manipulate it in specific situations ^1–5^. Long-term semantic representation is essential because it allows us to generalise knowledge gained from previous experience into novel situations ^6^. If we have a stored representation of the typical characteristics of, for example, a particular breed of dog, when we meet a new member of that breed we can predict its likely behaviour. Cognitive control over the activation and selection of semantic information is also critical because we store a wide range of information about any particular concept and different aspects of this knowledge are relevant in different circumstances ^7–9^. When playing a piano, for example, the functions of the keys and pedals are of central importance. But if one is asked to move the piano across the room, this dominant knowledge is not relevant and must be ignored in favour of focusing on the size, weight and value of the instrument.

There is substantial evidence that the *representation* and *control* elements of semantic cognition are supported by different neural networks and can be independently impaired as a consequence of brain damage ^10–13^. Little is known, however, about the relationships between these abilities in the healthy population. Do people who know more have better control of their knowledge, or is the quantity of semantic knowledge an individual possesses independent of their ability to regulate this information? Is cognitive control over semantic information strongly correlated with executive control ability in other domains? And do semantic representation and control decline in parallel as a function of healthy ageing? The present study was designed to investigate these questions.

Evidence from multiple methodologies indicates that semantic representation and control rely on distinct (but interacting) brain regions. The anterior temporal cortices act as a store of conceptual representations, through interaction with modality-specific association regions ^14–18^. Damage to this region results in a profound loss of semantic knowledge, as seen in the neuropsychological syndrome of semantic dementia ^16,19^. In contrast, the controlled use of semantic knowledge is associated with activation in a network comprising inferior prefrontal cortex, posterior middle temporal gyrus and the intraparietal sulcus ^9,20,21^. Damage to these regions following left-hemisphere stroke results in impaired ability to inhibit irrelevant semantic information, to retrieve weak or less automatic semantic associations and to understand ambiguous words with competing meanings ^10,22,23^.

The brain regions activated by tasks with high semantic control demands overlap partially with those implicated in the “multiple demand” network, which respond to increased control demands across many cognitive domains ^24,25^. This suggests that processes involved in control of semantic knowledge may overlap with those involved in executive control in other domains. In fact, fMRI studies suggest that semantic control can be fractionated into semantic-specific and domain-general components, which activate neighbouring areas of left inferior prefrontal cortex ^26,27^. The first of these is a <u>controlled retrieval</u> mechanism that is engaged when automatic activation of semantic knowledge is insufficient to complete a given task. For example, a participant completing a semantic association task might be asked whether *bee* is associated with *flower* or *tree*. In this situation, automatic processing of the cue may result in activation of the strongest associates of bees (*honey, hive* etc.) but fail to activate either of the available response options. Under these circumstances, individuals are assumed to engage in a goal-directed controlled search through the semantic store for the relevant information ^28^. This controlled retrieval process appears to be specific to semantic knowledge. In functional neuroimaging studies, increased need for controlled retrieval is associated with increased activation in the anterior, ventral portion of left inferior prefrontal cortex (BA47) ^26,29–31^. Structural and functional connectivity studies indicate that BA47 is strongly connected with the anterior temporal regions that code semantic representations, supporting the view that it is linked specifically with the regulation of semantic knowledge ^32–34^.

The second mechanism, here termed <u>semantic selection</u>, is engaged when automatic activation of semantic knowledge results in competition between multiple competing representations, which must be resolved. This is commonly assessed using the feature association task, in which participants are required to match concepts based on specific shared properties ^9^. For example, a participant might be asked whether a *wasp* or a *hive* is the same size as a *bee*. In this case, both response options are semantically associated with the cue and they compete with one another for selection. Top-down selection processes are thought to resolve the competition, based on the requirements of the current task ^28,35^. This may involve inhibition of pre-existing but irrelevant semantic links (e.g., *bee-hive*). Of course, the need to select from competing responses is not specific to semantic processing and is a major element of tests of inhibitory executive function like the Stroop task 36. Increased selection demands in semantic tasks are associated with activation in the posterior portion of left inferior prefrontal cortex (BA44/45) ^9,26,30^. This area is also activated by conditions of high competition in working memory tasks ^37^, the Stroop task ^38^ and other inhibitory tasks ^39^ and it shows strong structural connectivity with the domain-general multiple demand network ^33^. These findings suggest that semantic selection may be served by a domain-general executive selection system.

To summarise, neuropsychological and neuroimaging studies indicate a double dissociation between representations of semantic knowledge and the processes that control how this knowledge is used. Semantic control processes also appear to be multidimensional, consisting of semantic-specific processes supporting controlled retrieval of information from memory as well as domain-general selection mechanisms. However, the relationship between these abilities in the healthy population is not clear. Put simply, if you know a lot, are you better or worse at regulating how you access and use this knowledge? Three different potential answers to this question are suggested by three different literatures. First, as the evidence from neuroimaging and neuropsychology suggests, the quantity of knowledge one has and the ability to use it in a controlled fashion may be largely independent of one another. Alternatively, decades of intelligence research suggests that much individual variation across a wide range of cognitive tasks can be attributed a single general factor (g) ^40–42^. Applied to the semantic domain, this would imply a positive relationship between representation and control: individuals with high general ability should perform well on tests of both. Finally, findings in the episodic memory literature raise the possibility of a negative relationship between quantity of semantic knowledge and ability to control access to this information. In episodic memory studies, when participants learn to associate a large set of information with one particular stimulus, they find it harder to retrieve any particular piece of information from the set (the fan effect; ^43,44^). Extended to the semantic domain, this effect could suggest that individuals with a richer store of knowledge experience greater interference between competing representations, placing greater demands on control processes. Relatedly, recent computational models suggest that word retrieval and recognition become less efficient as individuals learn more words, because lexical items become harder to discriminate from one another ^45,46^. Although these models do not address the question of executive control directly, they also suggest that acquisition of greater knowledge brings with it additional processing costs.

The relationship between semantic control and executive control in other cognitive domains is also unresolved. As discussed earlier, some researchers have suggested that semantic control can be fractionated into domain-general and semantic-specific components. However, no studies have directly contrasted semantic control ability with non-semantic tests of executive function in healthy individuals. Such comparisons have been conducted in semantically-impaired stroke patients, who show a positive correlation between semantic control ability and performance on tests of non-semantic executive function ^10,47^. These findings could be taken as evidence that controlled processing of semantic information shares executive resources with other cognitive domains. However, as stroke patients typically have large lesions that span multiple frontal and parietal sites, it may simply reflect greater impairment in multiple, independent systems in patients with larger lesions. The present study provides a new perspective on this debate by assessing the relationship between semantic control and general executive ability in healthy individuals for the first time.

The final issue addressed in this study is the effect of healthy ageing on semantic abilities. It is typically the case that scores on semantic tests remain relatively stable across adulthood, in contrast to the marked declines seen in many other cognitive abilities ^42,48–53^. Most large-scale studies of cognitive aging have employed vocabulary tests as a measure of semantic ability. Such tests typically require participants to define words or perform multiple-choice judgements on the synonyms or antonyms of words. Meta-analysis indicates that older adults (over 60 years) actually score substantially higher on such tests than young adults below 30 years; ^49^. Such differences are typically explained in terms of older participants continuing to add to the semantic knowledge store throughout their lives. Indeed, greater time spent in formal education appears to account for much of the age effect ^49^. Vocabulary tests, which typically probe knowledge of low frequency words, provide a good indication of the breadth and richness of an individual’s semantic knowledge store. However, they are not designed to probe executively demanding aspects of semantic processing, such as controlled retrieval of concepts or competition resolution. As a consequence, the effect of ageing on semantic control abilities has not been assessed. This is important because it is possible that this aspect of semantic processing suffers a different fate to that seen for vocabulary measures. Some forms of executive function, outside the semantic domain, deteriorate as people grow older ^54–57^. Furthermore, a recent meta-analysis of functional neuroimaging studies indicates that older people show less activation in brain regions associated with semantic control during semantic processing ^58^. It is therefore possible that controlled use of semantic information shows age-related decline, particularly if this shares neural resources with other executive functions. Such declines may be evident even if the amount of knowledge in the semantic system increases.

The present study was designed to investigate differences in semantic abilities in young and older adults, as well as relationships between semantic knowledge, semantic control and non-semantic executive function in each age group. This was achieved by administering tests that probed the size of the semantic knowledge store (i.e., measures of vocabulary size) as well as tests that placed greater demands on controlled use of semantic knowledge (controlled retrieval and semantic selection). We assessed controlled semantic retrieval and semantic selection with separate tests, since these two elements of semantic control appear to dissociate neurally. Of course, these elements of semantic cognition are highly interactive and we assumed that all semantic tasks require each of them to some degree. Thus, none of the tests were expected to index a single element exclusively but they were designed to place greater demands on one element or another. We expected to replicate the established finding that older people have larger and richer repositories of semantic knowledge. However, it was predicted that performance on tests of semantic control would decline in old age.

## Method

### Participants

Fifty young adults, aged between 18 and 30, were recruited from the undergraduate Psychology course at the University of Edinburgh and participated in the study in exchange for course credit. Fifty older adults, aged between 61 and 91, were recruited from the Psychology department’s volunteer panel. A wide range of ages across later life were sampled, to allow for investigation of age effects within the older group as well as comparisons between young and older people. All participants reported to be in good health with no history of neurological or psychiatric illness. Demographic information for each group is shown in Table 2. Young and older adults did not differ significantly in years of education completed (*t*(98) = 0.49, *p* = 0.63). Proportions of male and females did not differ significantly between groups (χ2=3.48, *p* = 0.06), although the proportion of females was somewhat higher in the young group. Informed consent was obtained from all participants and the research was performed in accordance with all relevant guidelines/regulations. The study was approved by the University of Edinburgh Psychology Research Ethics Committee.

**Table 2:** *Demographic information and mean test scores for young and older participants*

| | Young adults | Older adults | Age $r$ in older group |
| --- | --- | --- | --- |
| $N$ | 50 | 50 | |
| Age | 19.1 (1.8) | 77.5 (8.6)*** |  |
| Sex M:F | 13:37 | 23:27 |  |
| Years of education | 13.8 (0.9) | 14.0 (3.1) | 0.11 |
| MMSE /30 | 28.8 (0.9) | 29.0 (1.2) | -0.28* |
| WCST (% correct) | 79.4 (9.3) | 56.2 (18.1)*** | -0.10 |
| WCST (categories) | 3.9 (1.2) | 1.7 (1.4)*** | -0.26 |
| Animals fluency | 25.4 (6.1)** | 21.8 (6.3) | -0.32* |
| Household objects fluency | 23.8 (6.5)* | 21.4 (5.1) | -0.28* |
| Letter F fluency | 13.8 (4.8) | 16.1 (6.5)* | -0.30* |
| Trails A errors | 0.1 (0.2) | 0.1 (0.3) | 0.04 |
| Trails B errors | 0.4 (0.8) | 0.7 (1.1) | 0.37** |
| Trails A time (s) | 27.8 (9.7) | 36.7 (11.7)*** | 0.64*** |
| Trails B time (s) | 46.8 (13.5) | 74.6 (31.3)*** | 0.55*** |
| Eriksen flanker task – Congruent trials accuracy | 0.98 (.02) | 0.99 (.02) | -0.24 |
| Eriksen flanker task – Incongruent trials accuracy | 0.96 (.04) | 0.98 (.03)** | -0.11 |
| Eriksen flanker task – Congruent trials RT (ms) | 578 (101) | 866 (263)*** | 0.33* |
| Eriksen flanker task – Incongruent trials RT (ms) | 640 (106) | 898 (252)*** | 0.35* |

*Standard deviations are shown in parentheses. Asterisks indicate the significance of $t$ -tests comparing young and older adults (Older adults column) and the Pearson's correlation between test scores and age within the older group ( $r$ column). \* = $p < 0.05$ ; \*\* = $p < 0.01$ ; \*\*\* = $p < 0.001$ .*

### General cognitive assessments

Participants completed a series of tests of general cognitive function and executive ability. The Mini-Mental State Examination was used as a general cognitive screen. All participants scored 25/30 or above and all were above the 10^th^ percentile for their age group and sex, according to UK normative data ^59^. Executive control was assessed using the Trail-making task ^60^. Executive function was also assessed with the Eriksen flanker task ^61^. Participants were presented with a row of five letters and pressed a key to indicate whether the central letter was a H or an N. They were instructed to ignore the four flanking letters on either side of the central target. On Congruent trials, the flanking letters matched the target (e.g., HHHHH) while on Incongruent trials they were inconsistent (e.g., NNHNN). The Incongruent trials therefore required participants to resolve competition between the two possible response options. As a test of complex executive function, a subset of participants (27 young; 26 older) completed a computerised version of the Wisconsin Card-Sorting Test (WCST), consisting of 64 trials ^62^. Finally, three categories of verbal fluency were administered, in which participants were given one minute to produce as many words as possible that fit a specific criterion. The criteria included two semantic categories (animals and household objects) and one letter of the alphabet (words beginning with F). Verbal fluency tasks require access to verbal-semantic knowledge but are also assumed to place high demand on executive function.

### Tests of semantic control

Participants completed a task that probed two forms of semantic control, using the same design as Badre et al. ^26^. The experiment consisted of 8 blocks of 12 trials. In four blocks, participants were asked to make semantic decisions based on global semantic association. Participants were presented with a probe word and asked to select its semantic associate from either two or four alternatives. The associate was either strongly associated with the probe (e.g., *town-city*) or more weakly associated (e.g., *iron-ring*; see Figure 1a). The 24 weak association trials were hypothesised to place greater demands on controlled retrieval of semantic information, because automatic spreading of activation in the semantic network would not be sufficient to identify the correct response ^28^.

**Figure 1:**
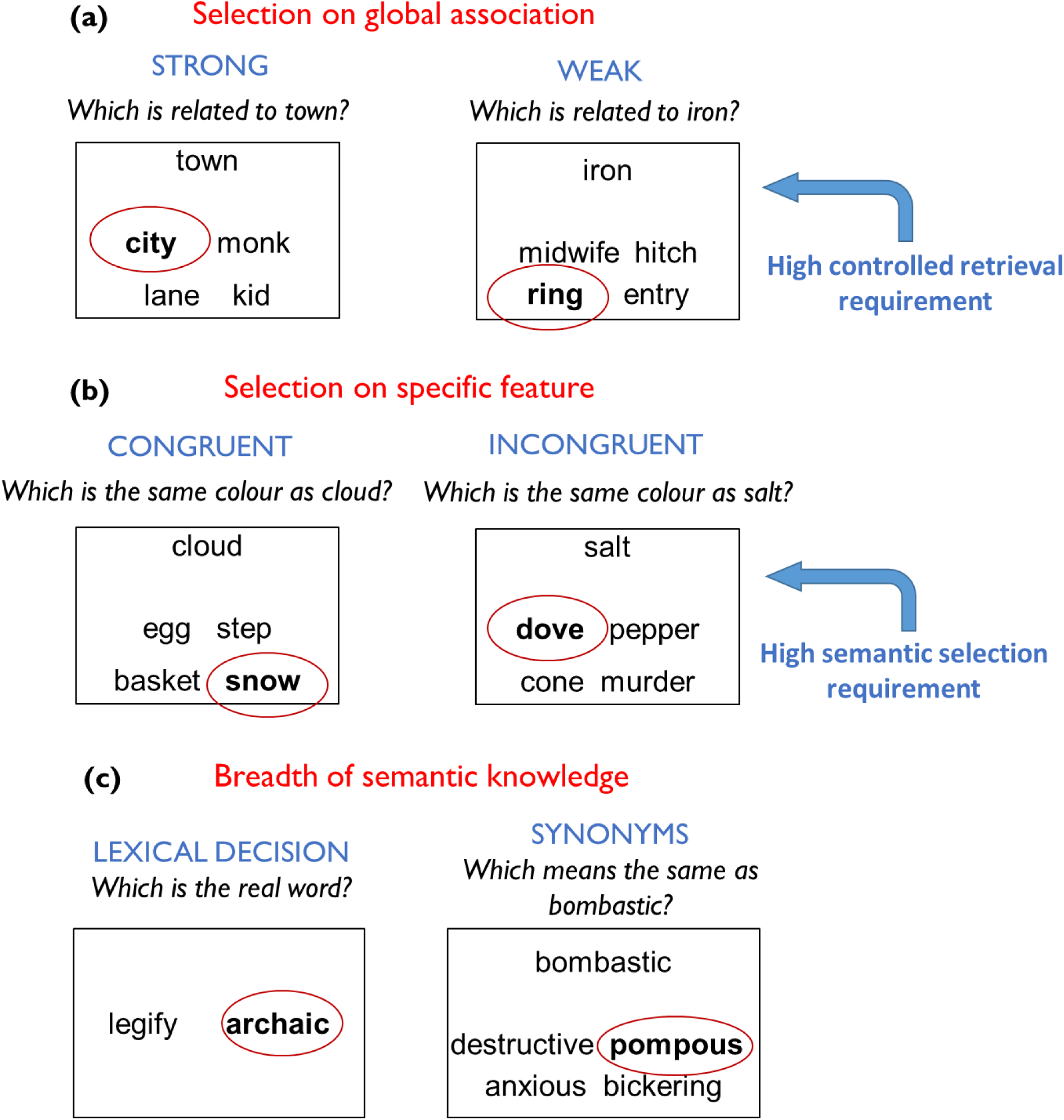
Example trials from (a) global association trials, (b) feature association trials and (c) tests of breadth of semantic knowledge. *The correct response is highlighted in each case*.

In the other four blocks, participants were asked to select items that matched on particular features (either Colour or Size). This task places high demands on semantic control because participants must direct attention away from global semantic associations and instead attend to specific item properties ^9^. At the beginning of each block, participants were given a feature to attend to (e.g., Colour). On each trial, they were provided with a probe and were asked to select, from two or four alternatives, the item that was most similar on the specified feature. There were 24 congruent trials, in which the probe and target shared a pre-existing semantic relationship, in addition to matching on the currently relevant feature (e.g., *cloud-snow*; see Figure 1b). On these trials, the foils did not match on the specified feature, nor did they share a global semantic association with the target. These trials therefore placed minimal demands on semantic selection mechanisms because the correct option was the one with the strongest pre-existing relationship with the probe. In contrast, on the 24 incongruent trials the probe and target shared no meaningful relationship, other than matching on the specified feature (e.g., *salt-dove*). Furthermore, one of the foils *did* have a strong semantic relationship with the probe, although it did not match on the currently relevant feature (*salt-pepper*). These trials placed high demands on selection mechanisms for two reasons: first, because there was no pre-existing semantic relationship between probe and target and second, because the strong but irrelevant relationship between the probe and foil had to be ignored.

Psycholinguistic properties of the words used in each condition are presented in Table 1. In the global task, words used in the strong and weak association conditions did not differ in mean frequency, concreteness or age of acquisition (all *p* > 0.48). Similarly, words used in the congruent and incongruent conditions of the feature association task did not differ in mean frequency, concreteness or age of acquisition (all *p* > 0.49). To quantify the strength of semantic relationships, distributed representations of word meanings were obtained from the word2vec neural network, trained on the 100 billion word Google News dataset ^63^. In common with other distributional models of word meaning, including latent semantic analysis ^64^, the word2vec model represents words as high-dimensional vectors, where similarity in two words’ vectors indicates that they appear in similar contexts, and thus are assumed to have related meanings. The word2vec vectors were used here as a recent study has shown that these outperform other available vector datasets in predicting human semantic judgements ^65^. The strength of the semantic relationship between two words was defined as the cosine similarity of their word2vec vectors. This value was calculated for the probe-target pairs in each condition. As expected, pairs in the global strong association condition were more strongly related than those in the weak association condition (*t*(46) = 4.55, *p* < 0.001). In the feature association task, the semantic relatedness of probe and target was stronger in the congruent condition (*t*(46) = 11.45, *p* < 0.001). The relationship between probes and foils was assessed in the same way. In the global association task these values were low in both conditions, indicating that the foils were semantically unrelated to the probes. This indicates that the foils in this task did not act as strong semantic competitors. However, in the feature association task, the relationship between probes and foils was significantly stronger in the incongruent condition (*t*(94) = 4.76, *p* < 0.001), reflecting the fact that a semantically related foil was present in this condition.

**Table 1:** *Mean psycholinguistic properties of words used in semantic tasks*

|  | Strong<br>Global | Weak<br>Global | Cong<br>Feature | Incon<br>Feature | Lex Dec | Synonyms |
| --- | --- | --- | --- | --- | --- | --- |
| Frequency (Zipf) | 4.23<br>(0.72) | 4.27<br>(0.81) | 4.28<br>(0.57) | 4.22<br>(0.65) | 2.49<br>(0.93) | 2.88<br>(1.14) |
| Concreteness | 4.40<br>(0.76) | 4.32<br>(0.73) | 4.68<br>(0.45) | 4.66<br>(0.41) | 3.76<br>(0.92) | 2.59<br>(0.83) |
| Age of acquisition | 6.17<br>(2.42) | 6.30<br>(2.12) | 5.97<br>(2.29) | 6.17<br>(1.99) | 11.97<br>(2.68) | 10.89<br>(2.97) |
| Probe-target<br>semantic<br>relatedness | 0.46<br>(0.16) | 0.27<br>(0.14) | 0.52<br>(0.13) | 0.15<br>(0.10) | - | - |
| Probe-foil<br>semantic<br>relatedness | 0.12<br>(0.08) | 0.09<br>(0.09) | 0.09<br>(0.07) | 0.23<br>(0.19) | - | - |

*Standard deviations are shown in parentheses. Word frequencies (on the log-transformed Zipf scale) were obtained from the SUBTLEX-UK database <sup>66</sup>. Concreteness values were obtained from Brysbaert et al. <sup>67</sup> and age of acquisition values from Kuperman et al. <sup>68</sup>. Semantic relatedness values were computed using distributed semantic vectors (see text for details). Cong = congruent; Incon = incongruent; Lex Dec = lexical decision.*

### Tests of breadth of semantic knowledge

Participants completed two tasks designed to probe the size of their store of semantic representations. These tasks probed knowledge of the meanings and identities of unusual words that were expected to be unknown to some members of the population. They therefore indexed the breadth of semantic knowledge available to each individual. The first task (Lexical Decision) was the Spot-the-Word Test from the Speed and Capacity of Language Processing battery ^69^. This was a lexical decision test comprising 60 pairs of letter strings (see Figure 1c). On each trial, participants were presented with a real word and a nonword and were asked to select the real word. Nonwords were phonologically and orthographically plausible, encouraging participants to rely on semantic knowledge for decision-making. The second task (Synonyms) was based on the Mill Hill vocabulary test ^70^, a multiple-choice test in which participants are asked to select the synonyms of particular words. There are two parallel forms of the test. To increase the difficulty of the task (thus avoiding ceiling effects), the hardest 22 trials from each parallel form were combined to make a new 44-item test. The test was presented in a four-alternative choice format (see Figure 1c).

To ensure that they placed greater demands on the semantic knowledge store, the words used in the semantic knowledge tests were significantly lower in frequency (*t*(662) = 20.7, *p* < 0.001) and concreteness (*t*(573) = 25.8, *p* < 0.001) and were later acquired (*t*(615) = 90.0, *p* < 0.001) than those used in the tests of semantic control (see Table 1).

### Procedure

Participants completed the general cognitive assessments followed by the semantic tasks. Semantic tasks were presented on a PC running Eprime 2.0 software. Participants first completed the Lexical Decision and Synonym tasks and then the semantic control experiment. Each task was preceded by a series of practice trials. Accuracy and reaction times (RTs) were recorded. Participants were instructed to respond as quickly as possible while avoiding mistakes. No time limit was placed on responses. They were encouraged to guess if unsure of the correct response.

### Statistical analyses

Reaction times were screened for outliers by winsorising any RTs more two standard deviations from a participant’s conditional mean (3.6% of trials). The results of the semantic control experiment were then analysed using mixed effects models to predict accuracy and RT at the level of individual trials. A 2 × 2 × 2 factorial model was first specified, including group as a between-subjects factor and task (global vs. feature judgments) and control demands (high vs. low) as within-subject factors. The weak association condition of the global judgements and the incongruent feature condition were considered to have high control demands. To investigate the two forms of semantic control independently, the two tasks were also analysed separately in 2 × 2 (control × group) models. Effects of age within the older group were also assessed, using a 2 × 2 (control × task) model that included age as a continuous predictor. To investigate performance on the semantic knowledge tasks, 2 × 2 mixed effects models were specified, including group as a between-subjects factor and task (lexical decision vs. synonyms) as a within-subjects factor. Finally, effects of age group on the individual conditions of the semantic control tasks and on the semantic knowledge tasks were assessed using t-tests. To control for multiple comparisons, a Holm-Bonferroni correction was applied to these results ^71^.

Mixed effects models were constructed and tested using the recommendations of Barr et al. ^72^. Linear models were specified for analyses of RT and logistic models for accuracy. We specified a maximal random effects structure for all models, including random intercepts for participants and items as well as random slopes for all predictors that varied within-participant or within-item. We also considered the following variables for inclusion in each model as covariates of no interest: trial position in test, number of response options, location of target, sex, education and MMSE score of participant. To avoid overfitting, a covariate was only included in a model if its inclusion significantly improved the fit of the model. The statistical significance of effects of interest was assessed by comparing the full model with a reduced model that was identical in every respect except for the exclusion of the effect of interest. Likelihood-ratio tests were used to determine whether the inclusion of the effect of interest significantly improved the fit of the model.

To explore the relationships between tests, Pearson’s correlations were computed within each age group separately. Test scores were also entered into principal components analysis, in which factors with eigenvalues greater than one were extracted and varimax rotated. These analyses were performed on accuracy data only, as RTs were likely to covary due to variations in general processing speed that were not specific to a particular cognitive domain. The following scores were included: Lexical Decision, Synonyms, Weak Associations, Incongruent Features, category fluency, letter fluency, WCST. Scores on low semantic control conditions and on the Trails and Erikson flanker tasks were not included due to ceiling effects. (These analyses were also repeated replacing Weak Association and Incongruent Features scores with the percentage effects of the manipulations of association strength and congruency. Results were very similar.)

All experimental materials and results are available on request from the author.

## Results

### General cognitive assessments

As shown in Table 2, young and older adults did not differ in MMSE scores. The fluency tasks showed varied effects of age, with young adults outperforming their older counterparts for the semantic categories while older people were significantly more successful at producing words beginning with F (although, within the older group, all scores were negatively correlated with age). Older people produced significantly fewer correct responses and completed fewer categories in the WCST. Completion times for the Trails task were analysed in a Part × Group ANOVA. Older adults were significantly slower overall (*F*(1,98) = 34.0, *p* < 0.001) and there was a significant interaction between part and group (*F*(1,98) = 22.9, *p* < 0.001), indicating that older adults showed greater slowing on Part B. This part has a greater executive demand because it requires switching between letters and numbers. RTs for the Eriksen flanker task were analysed using Condition × Group ANOVA. The older group were substantially slower (*F*(1,98) = 50.9, *p* < 0.001) but there was no interaction with condition (*F*(1,98) = 3.16, *p* = 0.08). Reaction times, but not accuracies, were correlated with age in the older group.

### Effects of age on semantic representation and control

Performance on the semantic control tasks are shown in Figure 2. These data were analysed in a 2 × 2 × 2 factorial mixed model that included group as a between-subjects factor and task (global vs. feature judgments) and control demands (high vs. low) as within-subject factors (where the weak associations and incongruent feature judgements were assumed to have high control demands). The results for accuracies and RT are shown in Table 3. Overall, older people were slower to respond but more accurate. As expected, the feature association task was more difficult than the global association task, with this effect being more pronounced in the older group (in accuracy only). The manipulations of control demand also had strong effects on accuracy and RT. Importantly, however, there were highly significant three-way interactions between these factors, suggesting that age had different effects on the two control manipulations.

**Figure 2:**
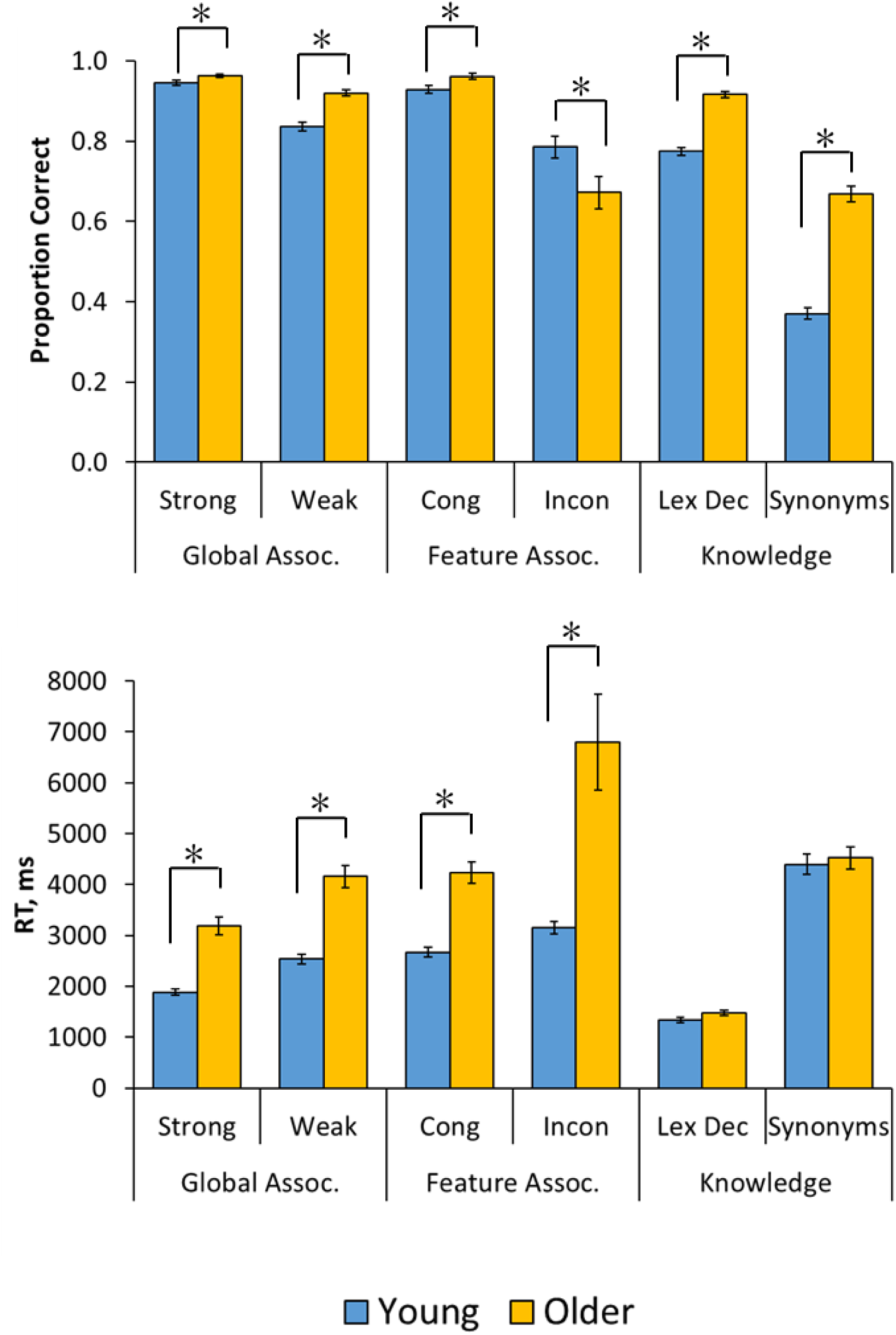
Performance on semantic tests. *Asterisks indicate significant age differences (Holm-Bonferroni-corrected p < 0.05). Cong = congruent; Incon = incongruent; Lex Dec = lexical decision*.

**Table 3:** *Analysis of semantic control experiment in young and older people*

| Effect | Accuracy |  | RT |  |
| --- | --- | --- | --- | --- |
|  | B (s.e.) | p | B (s.e.) | p |
| Group | .192 (.097) | .021 | .250 (.029) | <.001 |
| Task | -.483 (.119) | <.001 | .138 (.015) | <.001 |
| Task x Group | -.216 (.084) | .006 | .008 (.008) | .363 |
| Control | -.748 (.120) | <.001 | .130 (.014) | <.001 |
| Control x Group | -.110 (.088) | .160 | .012 (.008) | .128 |
| Task x Control | -.246 (.121) | .032 | -.008 (.014) | .594 |
| Task x Control x Group | -.193 (.090) | .022 | .024 (.008) | .002 |

The nature of the interaction was investigated with separate 2 × 2 (control × group) analyses for each task. In the global association task, there was a main effect of group for accuracy (B = .434, s.e. = .133, *p* < .001) but the effect of the control manipulation did not interact with group (B = .075, s.e. = .131, *p* = .56). For RT, there was again a main effect of group (B = .243, s.e. = .027, *p* < .001) but no interaction with control (B = -.011, s.e. = .010, *p* = .25). In other words, both groups showed similar effects of the control manipulation, despite older people being slower and more accurate overall. The picture was markedly different for the feature association task. Here, there was no overall difference in accuracy between young and older people (B = .003, s.e. = .123, *p* = .98). There was, however, a control × group interaction (B = -.346, s.e. = .121, *p* = .004) because the older group performed particularly poorly on the incongruent (high control) judgements relative to the young. Indeed, the incongruent feature judgements were the only condition in which older people made more errors than the young (see Figure 2). Analysis of reaction times indicated the older people were slower overall (B = .269, s.e. = .028, *p* < .001) and that there was again a control × group interaction (B = .037, s.e. = .012, *p* = .004) because they were particularly slow in the incongruent condition.

The picture that emerges from these results is that different forms of semantic control are affected differently by ageing. On the feature association task, older people demonstrated particular difficulty with semantic selection, when required to inhibit strong but irrelevant associations. In contrast, on the global association task the manipulation of association strength affected both groups to a similar degree, and older people performed more accurately overall. To investigate whether similar effects were present as a function of age within the older group, further analyses were performed in this group only, using models that included age as a continuous covariate (see Table 4). RT, but not accuracy, tended to decline as a function of age. Within the accuracy data there was, however, a trend towards an interaction between task, control and age, analogous to that seen in the between-group analysis (*p* = .088). Inspection of correlation coefficients indicated that this effect was driven by a negative association between age and performance that was only present on the incongruent trials of the feature association task (*r*(48) = −0.32, *p* = 0.024). Thus, within the older group there was a specific age-related decline in the ability to select task-relevant semantic relationships while ignoring irrelevant associations. No such interactions were observed for RT.

**Table 4:** *Analysis of effects of age within the older group for the semantic control experiment*

| Effect | Accuracy |  | RT |  |
| --- | --- | --- | --- | --- |
|  | B (s.e.) | *p* | B (s.e.) | *p* |
| Age | -.010 (.118) | .404 | .198 (.032) | <.001 |
| Task | -.704 (.162) | <.001 | .148 (.017) | <.001 |
| Task x Age | -.062 (.105) | .582 | .003 (.009) | .771 |
| Control | -.860 (.163) | <.001 | .144 (.017) | <.001 |
| Control x Age | -.179 (.112) | .094 | .009 (.009) | .343 |
| Task x Control | -.430 (.157) | .004 | .017 (.017) | .302 |
| Task x Control x Age | -.167 (.103) | .088 | .009 (.008) | .264 |

Performance on the two tests probing breadth of semantic knowledge is also shown in Figure 2. A 2 × 2 (task × group) mixed effects model indicated that older people performed much more accurately than young people on these tasks (B = .950, s.e. = .106, *p* < .001). Performance was generally more accurate on the lexical decision task (B = −1.396, s.e. = .175, *p* < .001), but this effect did not interact with group (B = -.007, s.e. = .078, *p* = .93). Analysis of RTs showed no group difference (B = .042, s.e. = .024, *p* = .084), which is surprising, given that the older group were significantly slower than young people across a range of other tasks. Thus, the accuracy and RT data both indicate that older people had a substantial processing advantage on these tasks, indicating that they had a larger and richer store of semantic representations available to them.

### Relationships among semantic and executive tests

A correlation matrix for test scores in the older group is presented in Table 5. Scores on the two semantic knowledge tests were strongly correlated, indicating that both indexed the breadth of participants’ semantic knowledge. Both of these tests were also positively correlated with years of education. In contrast, there were no correlation between the two tests of semantic control, suggesting that these tapped distinct abilities. However, both semantic control tasks showed weak positive correlations with one or both of the representation tests, suggesting a tendency for individuals with more developed semantic representations to be better at exercising control. Performance on the incongruent feature condition and on the verbal fluency tasks was negatively correlated with age.

**Table 5:** *Correlation matrix for task performance in older group*

|  | 1. | 2. | 3. | 4. | 5. | 6. | 7. | 8. | 9. |
| --- | --- | --- | --- | --- | --- | --- | --- | --- | --- |
| 1. Lexical Decision | - |  |  |  |  |  |  |  |  |
| 2. Synonyms | .82*** | - |  |  |  |  |  |  |  |
| 3. Weak Associations | .29* | .36* | - |  |  |  |  |  |  |
| 4. Incongruent Features | .35* | .19 | .16 | - |  |  |  |  |  |
| 5. Animals fluency | .21 | .20 | .32* | .34* | - |  |  |  |  |
| 6. HHO fluency | .13 | .18 | .30* | .09 | .55*** | - |  |  |  |
| 7. Letter fluency | .04 | .16 | .40** | .14 | .39** | .32* | - |  |  |
| 8. WCST | .05 | -.11 | -.09 | .15 | -.03 | .03 | -.38 | - |  |
| 9. Age | .05 | .13 | -.12 | -.32* | -.32* | -.28* | -.30* | -.10 | - |
| 10. Education | .39** | .42** | .15 | .08 | -.23 | -.12 | .05 | -.09 | .11 |

HHO = household objects. \* = $p < 0.05$ ; \*\* = $p < 0.01$ ; \*\*\* = $p < 0.001$ .

To explore the underlying commonalities among tests in more detail, a principal components analysis was performed. There were three factors with eigenvalues greater than one, which together accounted for 67% of the variance in scores (see Table 6). The weak association task and the verbal fluency tasks loaded on the first factor. In common with the weak association task, verbal fluency requires flexible, controlled retrieval of semantic knowledge. The lexical decision and synonyms tasks loaded strongly on the second factor, which appeared to index the breadth of participants’ semantic knowledge. The third factor had strong loadings for the WCST, suggesting that it captures aspects of domain-general executive function that are not specific to semantics. Interestingly, the incongruent feature condition also loaded most strongly on this factor, although it also had weaker loadings (>0.3) on the other two factors.

**Table 6:**
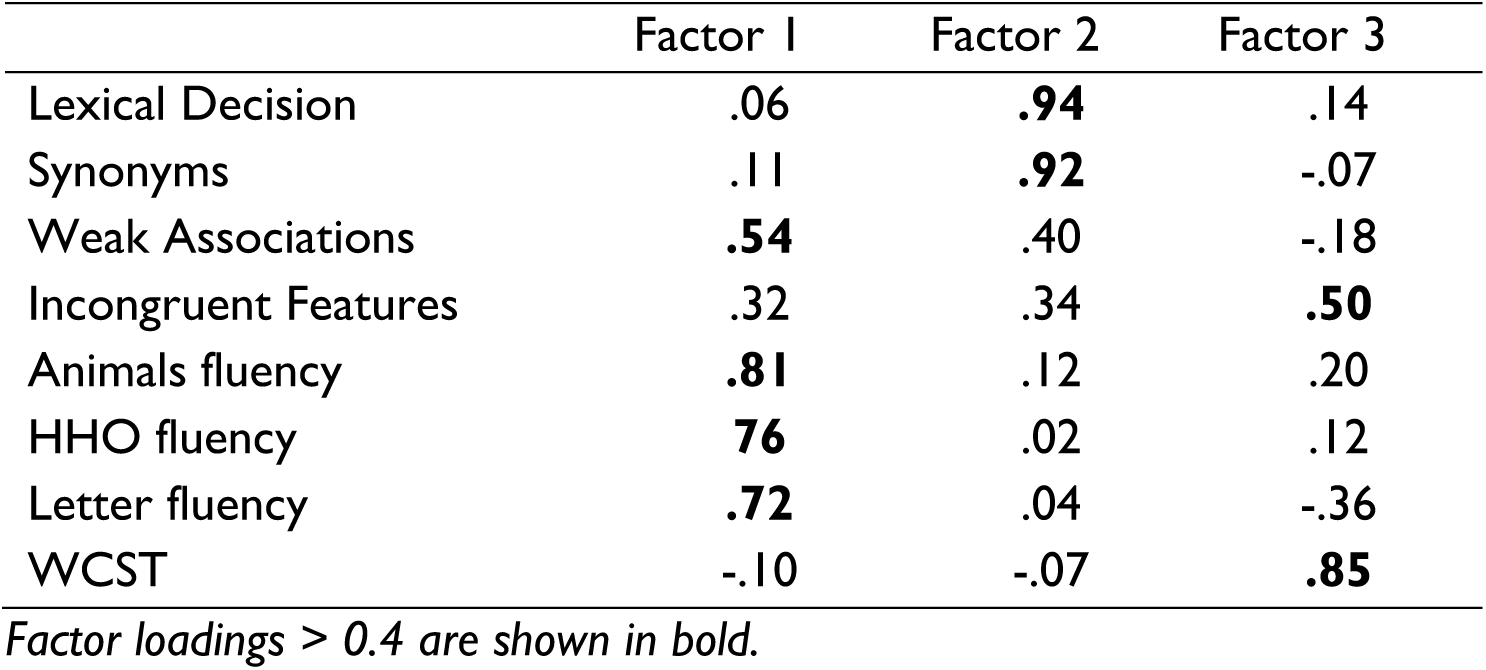
*Principal components analysis for the older group*

The correlation matrix for test scores in the young group is shown in Table 7 (note that correlations with age and educational level were not computed because there was little variation in these characteristics in the young group). As in the older group, there was a strong positive relationship between the two representation tests. There were also positive associations between these tests and the weak association task, although no correlation with the incongruent feature task. There was, however, a significant positive association between the incongruent feature task and the WCST.

**Table 7:** *Correlation matrix for task performance in young group*

|  | 1. | 2. | 3. | 4. | 5. | 6. | 7. |
| --- | --- | --- | --- | --- | --- | --- | --- |
| 1. Lexical Decision | - |  |  |  |  |  |  |
| 2. Synonyms | .64*** | - |  |  |  |  |  |
| 3. Weak Associations | .62*** | .29* | - |  |  |  |  |
| 4. Incongruent Features | .13 | .13 | .16 | - |  |  |  |
| 5. Animals fluency | .23 | .25 | .21 | .11 | - |  |  |
| 6. HHO fluency | -.10 | -.01 | .02 | -.16 | .45** | - |  |
| 7. Letter fluency | .20 | .10 | .17 | .12 | .42** | .24 | - |
| 8. WCST | .11 | .16 | .03 | .43* | .20 | .16 | .05 |

\* = $p < 0.05$ ; \*\* = $p < 0.01$ ; \*\*\* = $p < 0.001$ . Age and Education are not included for the young participants, as the range in these variables was too restricted to give valid results.

The results of the principal components analysis for the young group are shown in Table 8. There were three factors with eigenvalues greater than one, which together accounted for 64% of the total variance in test scores. The lexical decision and synonym tasks loaded strongly on the first factor but, unlike the older group, the weak association task also loaded strongly on this factor. The second factor had strong loadings for the verbal fluency tasks. However, the two semantic control tasks did not load on this factor. Finally, the third factor had strong loadings for the incongruent features and WCST, again suggesting the resolution of semantic competition is linked to domain-general executive resources.

**Table 8:** *Principal components analysis for the young group*

|  | Factor 1 | Factor 2 | Factor 3 |
| --- | --- | --- | --- |
| Lexical Decision | **.92** | .05 | .05 |
| Synonyms | **.74** | .08 | .12 |
| Weak Associations | **.76** | .11 | -.01 |
| Incongruent Features | .18 | -.10 | **.77** |
| Animals fluency | .23 | **.80** | .15 |
| HHO fluency | -.18 | **.81** | -.13 |
| Letter fluency | .20 | **.63** | .09 |
| WCST | -.06 | .18 | **.75** |

*Factor loadings > 0.4 are shown in bold.*

## Discussion

This study investigated relationships between semantic knowledge, semantic control and non-semantic executive control in young and older adults. Little is known about the relationships between these capabilities in healthy adults. However, there are clear neural dissociations between the representation of semantic knowledge and processes that control access to and use of this knowledge. Supporting proposals from cognitive neuroscience ^26,27^, the present study found evidence for two distinct forms of semantic control – controlled retrieval of knowledge and selection among competing representations. Performance on tests tapping these two abilities was independent of one another. In fact, while controlled retrieval ability was maintained in older people, there was a clear decrement in semantic selection ability, with older adults less able to ignore irrelevant semantic associations in favour of task-relevant aspects of knowledge. This was observed in comparisons of young and older adults and in correlational analyses of age effects within the older group. In both groups, semantic selection ability was linked with performance on a non-semantic executive function task, the Wisconsin card-sorting test, which also requires inhibition of irrelevant aspects of stimuli. In contrast to this specific decline in executively-demanding semantic tasks, older people had markedly larger stores of semantic knowledge, out-performing their younger counterparts on tests of knowledge for low frequency words. Taken together, these findings suggest three distinct contributors to semantic cognition: a store of representations which accumulates throughout the lifespan, processes for controlled retrieval of weakly related concepts from this store which appear to be age-invariant, and semantic selection mechanisms that decline in old age and are more closely associated with domain-general executive control.

Previous studies have observed that older people often perform better on semantic tests that probe vocabulary size ^42,48–50,52^. This effect was particularly striking in the present study: in addition to out-performing young people in accuracy, the older group matched them in reaction time, despite being slower to respond on a range of other tasks. These findings clearly indicate that older people have a more detailed store of knowledge representations that allow them to more successfully recognise and determine the meanings of low frequency words. In line with previous studies ^49^, older adults who had spent greater time in education exhibited broader semantic knowledge, indicating that educational experience plays an important part in determining the eventual size of the knowledge store. Consistent with this conclusion, a recent study found that the volume of the ventral temporal cortices in later life (a key neural substrate for the representation of semantic knowledge; e.g., ^16^) was a predictor of performance on semantic knowledge tests, and that this effect was mediated by educational level ^15^. It is important to note, however, that the young and older groups in the present study did not differ in mean years of education, so general life experience must also contribute to the large disparity in knowledge between the two groups.

The weak association task was used to probe participants’ ability to engage in controlled retrieval of less salient information from the semantic store. This aspect of control has been closely associated with anterior, ventral prefrontal cortex, which is thought to provide top-down constraint over the activation of knowledge represented in the temporal cortices ^26^. This ability to regulate retrieval of semantic information was closely related to breadth of semantic knowledge in young participants. Those with more detailed representations were better at detecting weak associations between concepts, which underscores the close interaction between these aspects of semantic cognition. The association of improved controlled retrieval with broader semantic knowledge, observed in both age groups, suggests that individuals with greater reserves of semantic knowledge also develop more effective mechanisms for regulating the retrieval of information from this store. In contrast, the retrieval of weak associations was not correlated with performance on the incongruent features task in either group. This suggests that there is a strong distinction between controlled retrieval of less salient knowledge and the ability to resolve semantic competition, despite both being considered aspects of semantic control. This conclusion is consistent with claims that these abilities have distinct neural correlates ^26,27^.

In contrast to their preserved ability to detect weak semantic associations, older adults showed a marked decline in semantic selection. This finding can be interpreted in the context of old-age deficits in inhibitory function and interference resolution across a range of tasks ^57,73,74^. Domain-general executive mechanisms may therefore be involved in this aspect of semantic processing, as implied by previous neuroimaging investigations ^28^. Supporting this conclusion, there were correlations between semantic selection ability and WCST performance in young people. The two tests also patterned together in principal components analysis in both groups, suggesting a common basis. An inhibitory account of these findings would hold that older people have difficulty suppressing strong pre-existing semantic associations in the incongruent condition of the feature association task, as well as inhibiting attention to irrelevant aspects of the stimuli in the WCST ^75^. Importantly, this account would not predict old-age decrements in the weak associations task, as there were no irrelevant associations to ignore in this task. It should be noted that results from the Erikson flanker task provided no evidence for impaired inhibitory function in older people, but this task has been criticised as a poor indicator of individual differences in inhibition ^76^.

In addition to domain-general inhibitory deficits, it is important to consider factors within the semantic system that might contribute to age-related decline in semantic selection. It is possible that the more extensive semantic knowledge of older participants exacerbates their problems with semantic selection, since they have a richer network of semantic associations that could potentially cause interference. This hypothesis makes two predictions. First, older people should perform more poorly than young people on tasks requiring selection among competing semantic representations, as was observed. Second, within each age group, those with broader semantic knowledge should display poorer semantic selection ability. There was no evidence for this second effect in the present study, which suggests that greater knowledge does not necessarily result in increased interference between concepts. An alternative possibility is that individuals with richer semantic knowledge develop more effective selection mechanisms to manage potential interference between concepts, as has been observed in bilinguals ^77^.

Finally, verbal fluency scores in the older participants were closely related to performance on the weak associations task, which indexes controlled retrieval of knowledge. In addition, in principal components analysis their scores on the fluency tasks loaded on the same factor as both semantic control tasks. Perhaps this is unsurprising, since verbal fluency tasks are widely assumed to tax the executive control system in addition to verbal knowledge reserves ^15,78–80^. Specifically, a strong ability to retrieve weak semantic associates is likely to boost the number of potential responses available in verbal fluency, while selection and inhibition processes are important for avoiding item repetition or out-of-category responses ^81^. There was no clear evidence for age-related declines in verbal fluency in the present study: category fluency scores were significantly lower in the older group but letter fluency scores were slightly higher. One possible explanation for this net stability is that the negative impact of impaired semantic selection in old age is offset by increased reserves of knowledge and improved controlled retrieval. More generally, the major contribution of this study is to indicate that semantic cognition relies on multiple distinct capabilities that vary substantially across individuals and age groups. This opens the possibility for future studies to explore the interplay between these elements of semantic cognition and their functional consequences for complex verbal and non-verbal semantic behaviours.

## Author contributions

The author devised the study, collected the data with assistance from student volunteers, analysed the data and wrote the paper.

## Competing financial interests

The author declares no competing financial interests.

## Acknowledgements

The work was undertaken by The University of Edinburgh Centre for Cognitive Ageing and Cognitive Epidemiology, part of the cross council Lifelong Health and Wellbeing Initiative (MR/K026992/1). Funding from the Biotechnology and Biological Sciences Research Council (BBSRC) and Medical Research Council (MRC) is gratefully acknowledged. I am grateful to Ellen Fearnley, Emily Hardy, Wing Yee Ho, Eszter Kalapos, Jasmine Kulay, Khushboo Mehra and Adam Ryde for assistance with data collection and to Beth Jefferies for sharing experimental stimuli.

